# *Staphylococcus warneri* dampens SUMOylation and promotes intestinal inflammation

**DOI:** 10.1101/2024.04.26.591263

**Authors:** Léa Loison, Marion Huré, Benjamin Lefranc, Jérôme Leprince, Moïse Coëffier, David Ribet

## Abstract

Gut bacteria play key roles in intestinal physiology, via the secretion of diversified bacterial effectors. Many of these effectors remodel the host proteome, either by altering transcription or by regulating protein post-translational modifications. SUMOylation, a ubiquitin-like post-translational modification playing key roles in intestinal physiology, is a target of gut bacteria. Mutualistic gut bacteria can promote SUMOylation, via the production of short- or branched-chain fatty acids (SCFA/BCFA). In contrast, several pathogenic bacteria were shown to dampen SUMOylation in order to promote infection. Here, we challenge this dichotomic vision by showing that *Staphylococcus warneri*, a non-pathogenic bacterium of the human gut microbiota, decreases SUMOylation in intestinal cells. We identified that Warnericin RK, a hemolytic toxin secreted by *S. warneri*, targets key components of the host SUMOylation machinery, leading to the loss of SUMO-conjugated proteins. We further demonstrate that the dampening of SUMOylation triggered by Warnericin RK promotes inflammation, and, more particularly, TNFα-dependent intestinal inflammatory responses.

Together, these results highlight the diversity of mechanisms used by non-pathogenic bacteria from the gut microbiota to manipulate host SUMOylation. They further highlight that changes in gut microbiota composition may impact intestinal inflammation, by changing the equilibrium between bacterial effectors promoting or dampening SUMOylation.

## Introduction

The gut microbiota plays essential roles in host physiology^1^. Metabolites produced by intestinal microorganisms are essential mediators of host-microbiota interactions^2^. They may regulate host cell activities by activating cell signalling pathways, by regulating cell transcription or by modulating host post-translational modifications^3^. Short-chain fatty acids (SCFAs) and branched-chain fatty acids (BCFAs), for example, were shown to participate to the maintenance of intestinal epithelial integrity and to dampen inflammation by promoting intestinal cell SUMOylation^4,5^.

SUMOylation is a post-translational modification consisting in the covalent addition of a small polypeptide, named SUMO (for <u>s</u>mall <u>u</u>biquitin-like <u>mo</u>difier), to target proteins^6,7^. The human genome codes for five SUMO paralogs, that share 45-97% sequence homology. SUMO1, SUMO2 and SUMO3 display ubiquitous expression patterns and are the most widely studied paralogs (SUMO4 and SUMO5 expression being restricted to only few, non-intestinal, tissues)^7^. SUMO2 and SUMO3 share 97% peptide sequence identity and are often referred to as SUMO2/3. SUMO1 and SUMO2/3 are not functionally redundant and are conjugated to distinct yet overlapping sets of proteins^8^. Conjugation of SUMO to target proteins occurs on lysine side-chains and requires an enzymatic machinery composed of an E1 enzyme (formed by the SAE1/SAE2 heterodimer), an E2 enzyme (Ubc9) and E3 enzymes^9^. SUMOylation is a reversible mechanism. Indeed, several proteases, called deSUMOylases, can cleave the covalent bond between SUMO peptides and their targets^10^. Conjugation of SUMO may have multiple consequences, including changes in protein activity, stability, localization, or interactions with other cellular components^6,7,11^. SUMOylation is more particularly involved in immunity and in intestinal homeostasis, as it limits detrimental inflammation and participates to tissue integrity maintenance^4,12–15^.

In contrast to mutualistic bacteria producing SCFAs and BCFAs, which increase intestinal SUMOylation, some pathogenic bacteria were shown to interfere with host SUMOylation in order to promote infection^16–18^. The food-borne pathogen *Listeria monocytogenes*, for example, secretes a pore-forming toxin called Listeriolysin O (LLO), which triggers a rapid degradation of the E2 SUMO enzyme, thereby inhibiting *de novo* SUMO-conjugations in host cells^19^. As deSUMOylases are still active in this case, this leads to a decrease in the level of SUMO-conjugated proteins in cells exposed to LLO, which has been correlated to an increase in bacterial infection^19,20^. Other pathogens were shown to target the host SUMOylation machinery such as *Shigella flexneri*, which induces a calpain-dependent cleavage of the E1 SUMO enzyme SAE2, or *Salmonella enterica* serovar Typhimurium, which downregulates the level of E2 SUMO enzyme using miRNA-dependent mechanisms^21,22^. *Staphylococcus aureus* was also shown to target the E2 SUMO enzyme in macrophages in order to promote intracellular survival^23^.

Our actual conception of the impact of gut bacteria on host SUMO-conjugation is thus dichotomic, with mutualistic bacteria on one hand that promote SUMOylation and pathogenic bacteria on the other hand which dampen SUMOylation. To challenge this concept, we focused on the gut bacterial species *Staphylococcus warneri*, a coagulase-negative staphylococci (CoNS) species naturally present in the human gut microbiota, but which was recently shown to be able to invade intestinal cells^24^. As such, this bacterium constitutes an intermediate situation between mutualistic bacteria and *bona fide* pathogens. In this study, we investigated the ability of *S. warneri* to interfere with SUMOylation and the potential associated consequences on intestinal physiology.

## Material and Methods

### Cell culture

Human Caco2 (American Type Culture Collection (ATCC)-HTB-37) and HeLa (ATCC-CCL2) cells were cultivated at 37°C in a 5% CO_2_ atmosphere in Minimum Essential Medium (MEM) supplemented with 10% fetal bovine serum (FBS), 2 mM L-glutamine (Invitrogen), non-essential aminoacids (Sigma-Aldrich), 1 mM sodium pyruvate (Gibco) and a mixture of penicillin (10000 U/mL) and streptomycin (10 mg/mL). Cell viability was quantified using the CellTiter-Glo^®^ Luminescent Cell Viability Assay (Promega).

### Bacterial strains

The bacterial strains used in this study were *Staphylococcus warneri* AW25 type strain (DSM 20316, DSMZ, Germany) and *Staphylococcus epidermidis* ATCC14990 type strain (DSM 20044, DSMZ, Germany). Staphylococcal strains were grown at 37°C in brain heart infusion (BHI) broth or on agar plates (BD Biosciences).

### Co-incubation of cells with bacterial supernatants and toxins

Warnericin RK (MQFITDLIKKAVDFFKGLFGNK) and delta-lysin II (MTADIISTIGDFVKWILDTVKKFTK) peptides, as well as their N-terminal formylated counterparts, were synthesized by Fmoc solid phase methodology on a Liberty microwave assisted automated peptide synthesizer (CEM, Saclay, France), using the standard manufacturer’s procedures at 0.1 mmol scale as previously described^25^. All Fmoc-amino acids (0.5 mmol, 5 eq.) were coupled on Fmoc-Lys(Boc)-HMP resin, by *in situ* activation with HBTU (0.5 mmol, 5 eq.) and DIEA (1 mmol, 10 eq.) before Fmoc removal with a 20% piperidine in DMF. After completion of the chain assembly, formylation of the N-terminal extremity was carried out on solid support, as previously described^26^. Briefly, the formylating agent was first prepared by mixing formic acid (25 mmol, 500 eq.) and diisopropylcarbodiimide (12.5 mmol, 250 eq.) in diethyl ether (14.5 mL) for 4h at 0°C. Then, after filtration and concentration under vaccum, the obtained solution (2 mL) was added to the peptidyl-resin (0.05 mmol) in 1 mL of DMF with DIEA (0.12 mmol, 2,4 eq.) and kept at 4°C overnight, followed by DMF and DCM washes of the resin. The two peptides and their formylated counterparts were deprotected and cleaved from the resin by adding 10 mL of the mixture TFA/TIS/H2O (9.5:0.25:0.25) for 180 min at room temperature. After filtration, crude peptides were washed twice by precipitation in TBME followed by centrifugation (4500 rpm, 15 min). The synthetic peptides were purified by reversed-phase HPLC (Gilson, Villiers le Bel, France) on a 21.2 x 250 mm Jupiter C18 (5 µm, 300 Å) column (Phenomenex, Le Pecq, France) using a linear gradient (50-90% over 45 min) of acetonitrile/TFA (99.9:0.1) at a flow rate of 10 mL/min. The purified peptides were then characterized by MALDI-TOF mass spectrometry on a ultrafleXtreme (Bruker, Strasbourg, France) in the reflector mode using α-cyano-4-hydroxycinnamic acid as a matrix. Analytical RP-HPLC, performed on a 4.6 x 250 mm Jupiter C18 (5 µm, 300 Å) column, indicated that the purity of all peptides was >99.9%. The net peptide content of lyophilized and dry peptide samples is assessed by CHN elemental analysis (N-content) using an Unicube elemental analyser (Elementar, Lyon, France). Briefly, samples were accurately weighed (∿ 200 µg) into tin boats and compressed with a manual pressing tool in order to remove remaining air from the packed sample and then combusted in pure oxygen under static conditions resulting in the formation of CO_2_, NOx, H_2_ and H_2_O. The NOx compounds were reduced to N_2_ using elemental cupper. Gases were separated by gas chromatography and detected quantitatively (acetanilide as calibrant) by a thermal conductivity detector.

Caco2 and HeLa cells were seeded at a density of 1.0 x 10^5^ cells/cm^2^ the day before co-incubations. *S. warneri* and *S. epidermidis* were cultured overnight at 37°C. Before co-incubations, cells were incubated in FBS- and antibiotics-free culture medium. Bacterial cultures were diluted in BHI to obtain a suspension with an optical density (OD) at 600 nm of 13.0. Supernatants were then collected after centrifugation of the diluted bacterial cultures 2 minutes at 19,000 x *g* at room temperature. Supernatants were filtered (0.2 µm filter, Sartorius), treated or not for one hour at 56°C with 0.4 mg/mL proteinase K (P8107S, New England Biolabs) and then heated or not for one hour at 85°C.

Supernatants and purified toxins were directly added in cell culture medium, as indicated in the text. After incubations, cells were washed twice with 1X PBS and lysed directly either in Laemmli buffer (4% SDS, 20% glycerol, 125 mM Tris-HCl [pH 6.8], 100 mM dithiothreitol [DTT], 0.02% bromophenol blue), for immunoblot analyses, or in RLT buffer (Qiagen), for RNA extraction.

### Immunoblot analysis

Cells lysates in Laemmli buffer were boiled for 5 minutes, sonicated and their protein contents were analysed by electrophoresis on TGX Stain-free pre-cast SDS-polyacrylamide gel (Bio-Rad). Proteins were then transferred on PVDF membranes (GE Healthcare) and detected after incubation with specific antibodies using ECL Clarity Western blotting Substrate (Bio-Rad). Mouse anti-actin (R5441, Sigma-Aldrich, 1:10,000 dilution), mouse anti-Ubc9 (610748, BD Biosciences, 1:1,000 dilution), rabbit anti-SAE1 (#13585, Cell Signaling Technology, 1:1,000 dilution), rabbit anti-SAE2 (D15C11, Cell Signaling Technology, 1:1,000 dilution) and rabbit anti-SUMO3 (R205 (is2)^27^, 1:5,000 dilution) antibodies were used as primary antibodies for immunoblotting experiments. HRP-conjugated goat anti-rabbit IgG (H+L) (BI 2407, Abliance, 1:5,000 dilution) and anti-mouse IgG (H+L) (BI 2413C, Abliance, 1:5,000 dilution) antibodies were used as secondary antibodies for immunoblotting experiments. Quantifications of proteins were performed on a ChemiDoc Imaging System (Bio-rad). Quantities of SUMO-conjugated proteins (above 50 kDa), SAE1, SAE2 and Ubc9 were normalized by actin levels.

### Quantification of proinflammatory cytokines expression

Cells incubated in FBS- and antibiotics-free culture medium were pre-treated with 10 µM TAK981 (Subasumstat, MedChemExpress) for 2 hours or with 20 µM Warnericin RK for 1 hour. 1 µg/mL Pam_3_CSK_4_ (InvivoGen) or 100 ng/mL recombinant human TNFα (PeproTech) were then added directly in cell culture medium. After 4 additional hours of incubation, cells were washed twice with 1x PBS and total RNAs were extracted using the RNeasy Plus Mini kit (Qiagen). 1 µg of total RNAs were reverse transcribed using random hexamers and M-MLV reverse transcriptase (Invitrogen). Quantification of gene expression was performed by qPCR using Itaq Universal SYBR Green Supermix (Bio-Rad). GAPDH was used as an internal reference for normalization. Primers used in this study are hGAPDH_F (5’-TGCCATCAATGACCCCTTCA-3’), hGAPDH_R (5’-TGACCTTGCCCACAGCCTTG-3’), hIL8_F (5’-TGGCAGCCTTCCTGATTT-3’), hIL8_R (5’-AACTTCTCCACAACCCTCTG-3’), hIL1β_F (5’-ACGAATCTCCGACCACCA-3’), hIL1β_R (5’-ATAAGCCTCGTTATCCCATG-3’), hCCL2_F (5’-GAAAGTCTCTGCCGCCCTTC-3’) and hCCL2_R (5’-ACAGATCTCCTTGGCCACAA-3’). Serial dilutions of target cDNAs were included on each plate to generate a relative curve and to integrate primer efficiency in the calculations of gene expression.

### Quantification of hemolytic activity

Hemolytic activities of *S. warneri* supernatant, Warnericin RK and delta-lysin II were determined by detecting the release of hemoglobin from sheep erythrocytes, as previously described^28^. Briefly, sheep erythrocytes washed in 1X PBS were incubated with purified toxins or dilutions of *S. warneri* supernatants for 30 minutes at 37°C and then centrifugated 5 minutes at 2,000 x *g* to pellet unlysed red blood cells. The OD at 540 nm of the resulting supernatants was then measured to estimate the percentage of lysed sheep erythrocytes. Erythrocytes hypotonically lysed in distilled water were used as positive controls (100% hemolysis).

## Results

### S. warneri inhibits host cell SUMOylation

To assess whether *S. warneri* regulates host cell SUMOylation, we first compared the global pattern of SUMO2/3-conjugated proteins in a human cell line (HeLa cells) incubated or not with *S. warneri* supernatant. We observed that *S. warneri* supernatant induces a significant decrease in the level of SUMO-conjugated proteins after 1h or 5h of incubation (Figure 1A and 1B). This decrease in SUMOylation is dependent on the amount of *S. warneri* supernatant added to cells (Figure 1).

**Figure 1:**
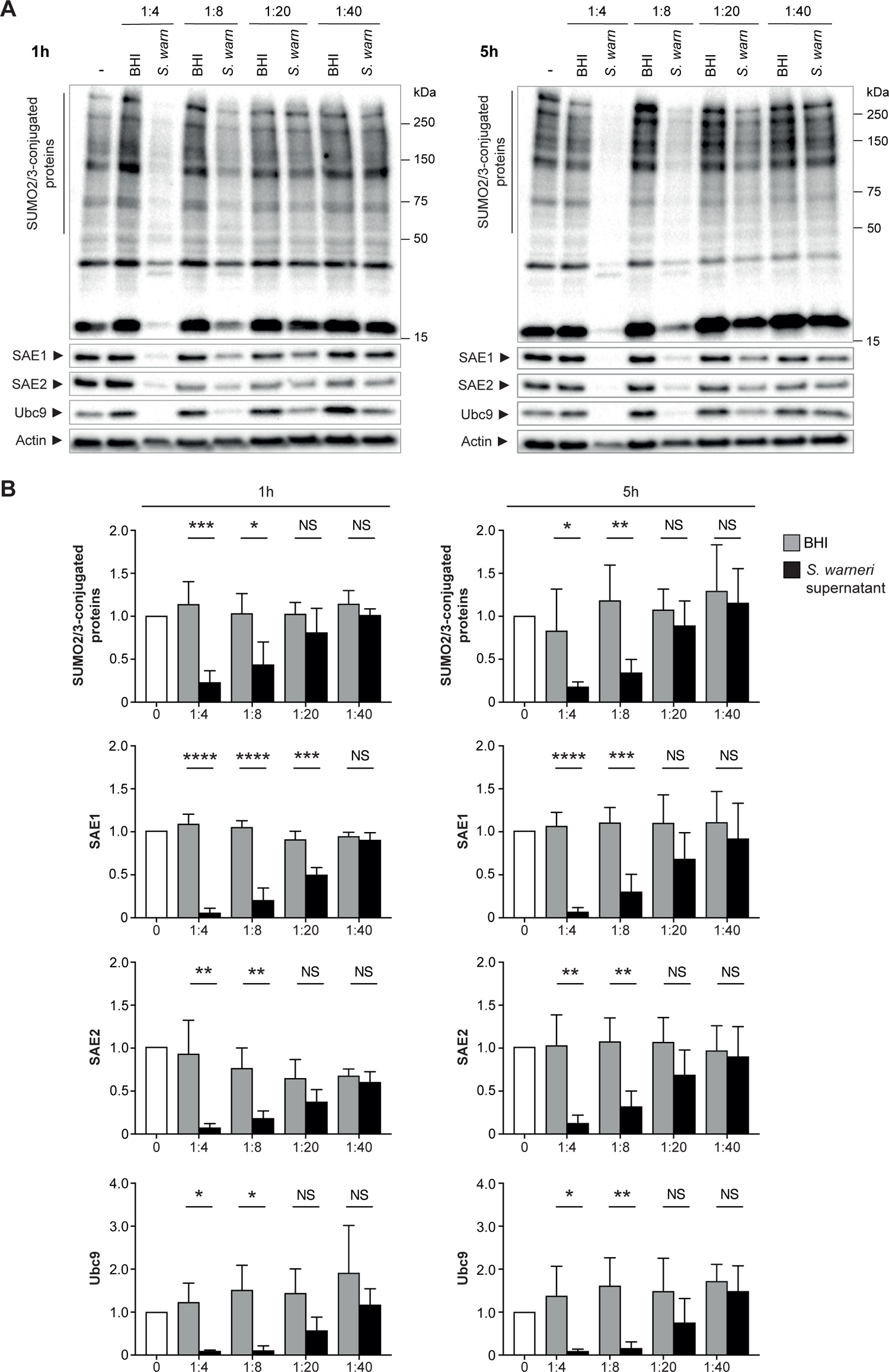
*Staphylococcus warneri* decreases SUMOylation in HeLa cells. A, Immunoblot analysis of SUMO2/3-conjugated proteins, SAE1, SAE2 and Ubc9 in HeLa cells treated with dilutions of *S. warneri* supernatant (*S. warn*) or with bacterial culture medium alone (BHI) for 1h (left) or 5h (right). B, Quantification of SUMO2/3-conjugated proteins (above 50 kDa), SAE1, SAE2 and Ubc9 levels after normalization by actin levels. Values are expressed as fold-change versus untreated cells (mean ± s.d.; n=3-4; *, *P*<0.05; **, *P*<0.01; ***, *P*<0.001; ****, *P*<0.0001; NS, not significant; two-tailed Student’s t-test).

As several host targets are deSUMOylated in response to *S. warneri* supernatant, we hypothesized that this bacterium directly impacts the host SUMOylation machinery, as already observed for pathogenic bacteria such as *L. monocytogenes*^19^. We thus quantified the levels of E1 and E2 enzymes in HeLa cells incubated with *S. warneri* supernatant and observed a significant decrease in both Ubc9, SAE1 and SAE2 levels, in a dilution-dependent manner (Figure 1A and 1B). Since SUMOylation levels correspond to a dynamic equilibrium between SUMO-conjugation and deSUMOylation reactions, this decrease in the level of SUMO-conjugating enzymes is likely responsible for the global decrease in SUMOylation triggered by *S. warneri*.

Together, these results suggest that *S. warneri* dampens host SUMOylation by secreting an effector targeting the human SUMO conjugation machinery. To determine whether the alteration of host SUMOylation triggered by *S. warneri* is specific to this bacterial species, we performed similar experiments with the supernatant of *Staphylococcus epidermidis,* another CoNS species. Interestingly, in contrast to *S. warneri*, we did not observe any changes in the global SUMOylation pattern of HeLa cells incubated with high (1:20) and low (1:8) dilutions of *S. epidermidis* supernatant for 5h (Figure 2A). The level of SUMO E1 and E2 enzymes remained unchanged in response to *S. epidermidis* (Figure 2B). These data suggests that the effector secreted by *S. warneri* and targeting host SUMOylation is absent from the supernatant of *S. epidermidis*.

**Figure 2:**
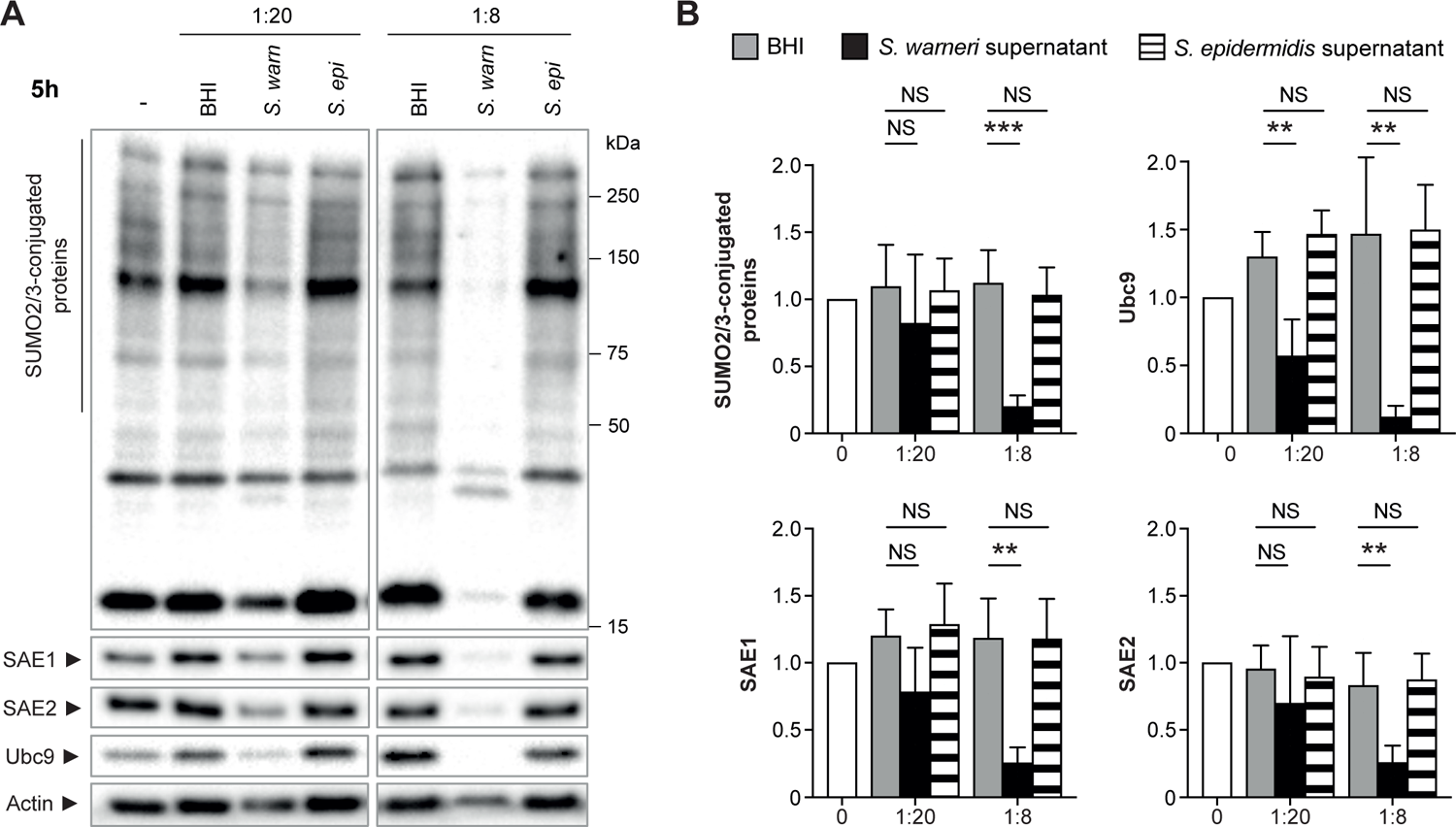
*S. epidermidis* does not inhibit host cell SUMOylation. A, Immunoblot analysis of SUMO2/3-conjugated proteins, SAE1, SAE2 and Ubc9 in HeLa cells treated with dilutions of *S. warneri* (*S. warn*) or *S. epidermidis* (*S. epi*) supernatants, or with bacterial culture medium alone (BHI), for 5h. B, Quantification of SUMO2/3-conjugated proteins (above 50 kDa), SAE1, SAE2 and Ubc9 levels after normalization by actin levels. Values are expressed as fold-change versus untreated cells (mean ± s.d.; n=4; **, *P*<0.01; ***, *P*<0.001; NS, not significant; One-way ANOVA, with Dunnett’s correction).

### Identification of the S. warneri effector triggering host protein deSUMOylation

In order to characterize the effector secreted by *S. warneri* and responsible for the observed decrease in host protein SUMOylation, we pre-treated *S. warneri* supernatant with proteinase K, a broad-spectrum serine protease. We also tested the effect of heat inactivation on *S. warneri* supernatant. We observed that heat inactivation did not impair *S. warneri*-induced deSUMOylations (Figure 3). In contrast, pre-treatment with proteinase K induces a complete blocking of *S. warneri*-induced deSUMOylations, as well as a lack of decrease in SUMO E1 and E2 levels (Figure 3). Together, these results suggest that the effector secreted by *S. warneri* and involved in host protein deSUMOylation is a heat-resistant protein or peptide.

**Figure 3:**
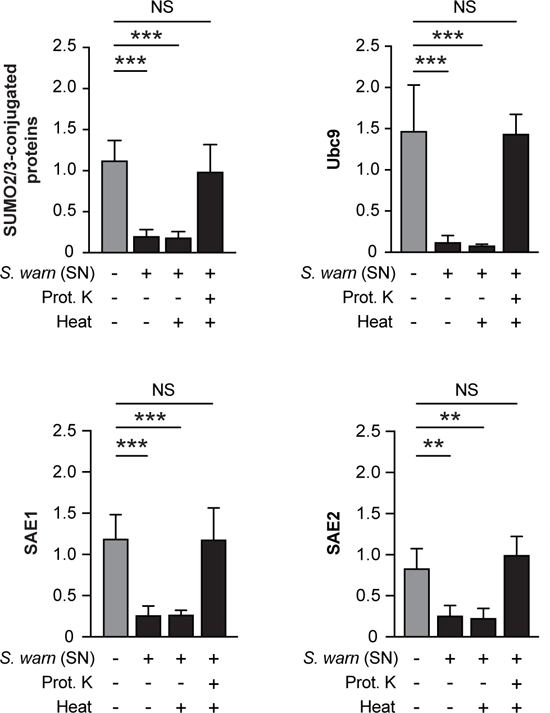
Characterization of the *S. warneri* effector interfering with SUMOylation. Quantification of SUMO2/3-conjugated proteins (above 50 kDa), SAE1, SAE2 and Ubc9 levels, after normalization by actin levels, in HeLa cells treated with *S. warneri* supernatant (SN) (1:8 dilution) or bacterial culture medium alone (BHI), with or without heat or Proteinase K (Prot. K) pre-treatments. Values are expressed as fold-change versus untreated cells (mean ± s.d.; n=4; **, *P*<0.01; ***, *P*<0.001; NS, not significant; One-way ANOVA, with Dunnett’s correction).

Since the effect of *S. warneri* on host SUMOylation is similar to the one described for *L. monocytogenes* or other bacterial pathogens secreting pore-forming toxins^17,19^, we focused on potential pore-forming proteins or peptides produced by *S. warneri,* which would not be encoded by *S. epidermidis*. Three peptides, named Warnericin RK (WRK), delta-lysin I (HldI) and delta-lysin II (HldII), were identified in various strains of *S. warneri* and were reported to exhibit hemolytic activities^29,30^. HldI and HldII belong to a family of peptides called delta-hemolysins (Hld), which are also encoded by *S. epidermidis* and other Staphylococcal species^30^. In contrast, WRK has no homolog in *S. epidermidis* (Figure 4)^29^. We thus tested the potential effect of WRK on host cell SUMOylation. We also tested, as a control, the effect of the HldII peptide (the gene coding for HldI being deleted in the *S. warneri* AW25 strain used in our experiments).

**Figure 4:**
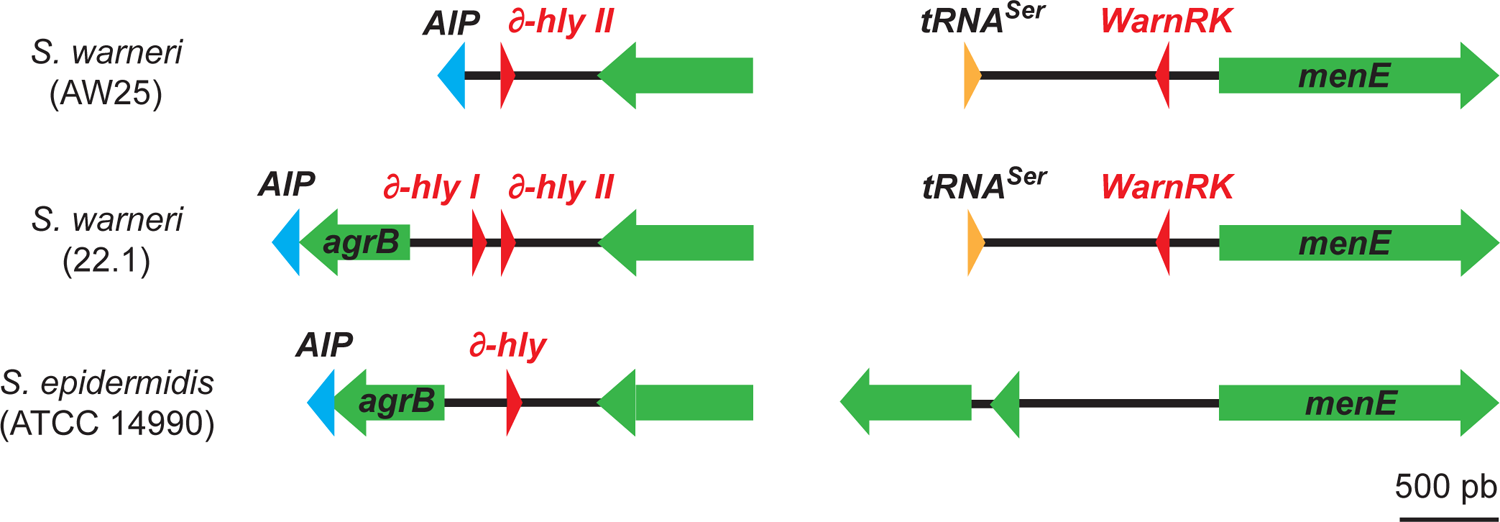
Genetic loci of hemolytic peptides encoded by *S. warneri* and *S. epidermidis* Schematic representation of the genetic loci coding for Warnericin RK (WarnRK), delta-lysin I (∂-*hlyI*) and delta-lysin II (∂-*hlyII*) in *S. warneri* AW25 type strain, *S. warneri* 22.1 strain and in *S. epidermidis* ATCC 14990 type strain.

Both Warnericin RK and HldII peptides were synthetized, either in their unmodified form or with an N-terminal formylation (formyl-WRK and formyl-HldII). Indeed, formylation of the N-terminal methionine residue has been reported for both peptides^29^. We first performed hemolytic assays to confirm the ability of these peptides to destabilize eukaryotic plasma membranes. We observed that both WRK and HldII peptides exhibit hemolytic activity and that the activity of WRK is significantly higher than the one of HldII (Figure 5A). Of note, N-formylation of WRK increases its hemolytic activity. We confirmed in parallel that the supernatant of *S. warneri* grown in stationary phase also exhibits hemolytic activity (Figure 5B).

**Figure 5:**
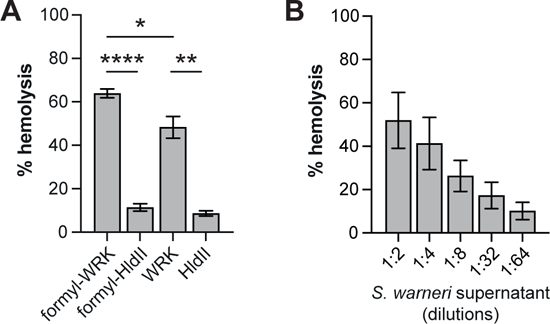
Hemolytic activities of Warnericin RK and delta-lysin II. A, Hemolytic activities of 25 µM WRK, formyl-WRK, HldII and formyl-HldII. Values are expressed as percentage of red blood cell lysis compared to a positive reference (100% lysis) (mean ± s.e.m; n=3; *, *P*<0.05, **, *P*<0.01; ****, *P*<0.0001; two-tailed Student’s t-test). B, Hemolytic activity of *S. warneri* supernatant (mean ± s.e.m; n = 3).

We then investigated the effect of Warnericin RK and delta-lysin II on HeLa cell SUMOylation. For this, we co-incubated cells with various concentrations of WRK and HldII peptides for 1 hour and analysed the level of SUMO E1 and E2 enzymes as well as the global pattern of SUMO2/3-conjugated proteins by immunoblotting experiments. Strikingly, we observed that WRK is sufficient to decrease the level of both SUMO E1 and E2 enzymes and to trigger a global dampening of host protein SUMOylation in a dose-dependent manner (Figure 6). Of note, the effect of the formylated form of WRK on host SUMOylation machinery is slightly stronger than the N-terminal free form (Figure 6). We performed in parallel viability assays and observed a limited toxicity of WRK at 20 µM on HeLa cells after 1 hour of incubation (< 30%) (Figure S1). In contrast to WRK, the HldII peptide does not significantly affect host SUMOylation, even at high concentrations (100 µM) (Figure S2). Together, these results demonstrate that Warnericin RK targets the host SUMOylation machinery and triggers a deSUMOylation of host proteins.

**Figure 6:**
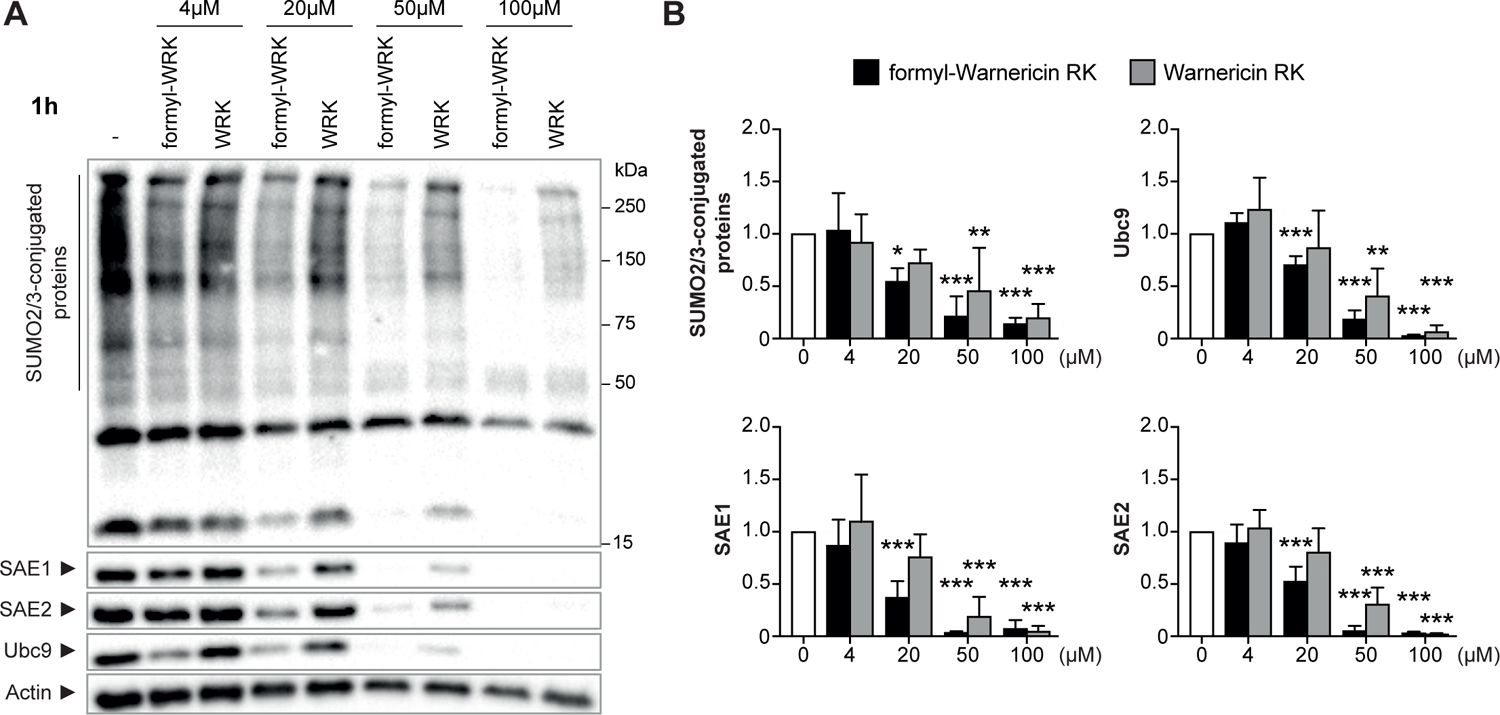
Warnericin RK decreases protein SUMOylation in HeLa cells. A, Immunoblot analysis of SUMO2/3-conjugated proteins, SAE1, SAE2 and Ubc9 in HeLa cells treated or not with formyl-WRK or WRK for 1h. B, Quantification of SUMO2/3-conjugated proteins (above 50 kDa), SAE1, SAE2 and Ubc9 levels, after normalization by actin levels. Values are expressed as fold-change versus untreated cells (mean ± s.d; n=4-5; *, *P*<0.05; **, *P*<0.01; ***, *P*<0.001; One-way ANOVA, with Dunnett’s correction).

### Warnericin RK affects intestinal SUMOylation

As *S. warneri* is a natural member of the human gut microbiota, we tested the effect of WRK on intestinal cell SUMOylation^24^. For this, Caco2 cells were incubated with 20 µM formyl-WRK for 1 and 5 hours and the level of SUMO E1 and E2 enzymes, as well as the global pattern of SUMO2/3-conjugated proteins, were then analysed by immunoblotting experiments (Figure 7). After five hours of incubation, we observed a significant decrease in the overall SUMOylation profile, as well as a decrease in the level of Ubc9, as previously observed in HeLa cells (Figure 7). Again, this decrease in SUMOylation is associated with a limited toxicity (<30%) (Fig. S1). Together, these results indicate that Warnericin RK promotes protein deSUMOylation in different human cell types, including intestinal cells, by targeting the E2 SUMO enzyme.

**Figure 7:**
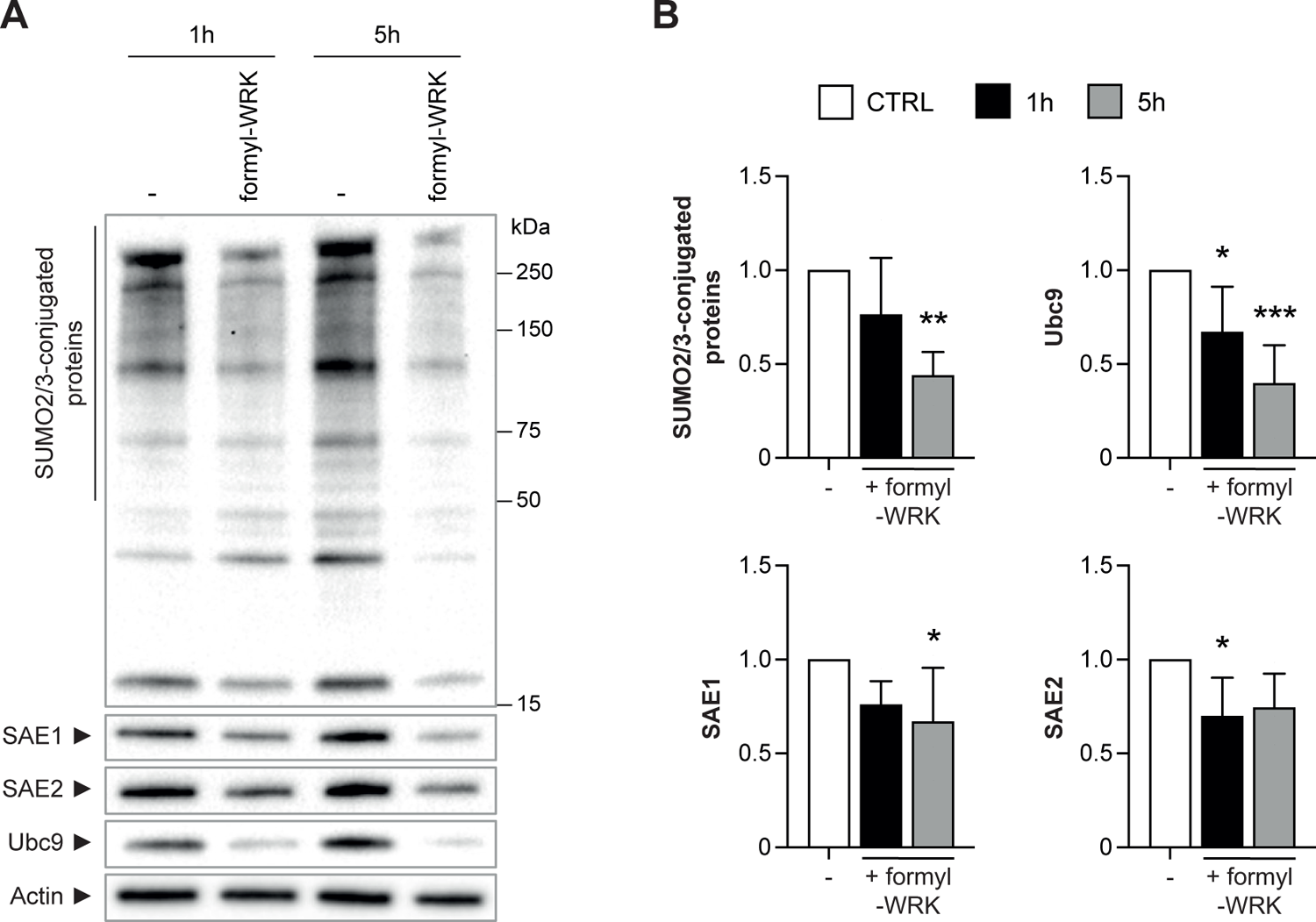
Warnericin RK induces deSUMOylation in intestinal cells. A, Immunoblot analysis of SUMO2/3-conjugated proteins, SAE1, SAE2 and Ubc9 in Caco2 cells treated with 20 µM formyl-WRK for 1h and 5h. B, Quantification of SUMO2/3-conjugated proteins (above 50 kDa), SAE1, SAE2 and Ubc9 levels, after normalization by actin levels. Values are expressed as fold-change versus untreated cells (mean ± s.d; n=4-5; *, *P*<0.05; **, *P*<0.01; ***, *P*<0.001; One-way ANOVA, with Dunnett’s correction).

### Warnericin RK-induced deSUMOylations promote inflammatory responses

As SUMOylation is known to regulate inflammation^13,14^, we investigated whether WRK-induced deSUMOylations modulate intestinal cells inflammatory responses. For this, Caco2 cells were first pre-treated or not with formyl-WRK for 1 h and then incubated for 4 h with recombinant human TNFα or Pam_3_CSK_4_, a synthetic triacylated lipopeptide acting as a ligand of Toll-like Receptor 1/2 (TLR1/2). After incubation, the expression levels of IL8, IL1ß and CCL2 cytokines in Caco2 cells were quantified by qRT-PCR. As a control, we treated cells with TAK981, a selective inhibitor of SUMOylation targeting the E1 SUMO enzymes^31^, to mimic WRK-induced deSUMOylations.

As expected, we observed that TNFα and Pam_3_CSK_4_ upregulates the expression of IL8, IL1ß and CCL2 in Caco2 cells. Interestingly, pre-treatment with formyl-WRK further increases IL8 upregulation in response to TNFα (Figure 8). Similar effects on IL8 were observed in cells pre-treated with TAK981, thus suggesting that the effect of WRK on IL8 expression is due to the deSUMOylations triggered by this bacterial toxin. In contrast to IL8, WRK has no effect on the upregulation of IL1ß and CCL2 expression in Caco2 cells treated with TNFα (Figure 8). The observed effects of WRK on cytokine expression appears to be specific to TNFα responses since WRK does not promote the expression of IL8, IL1ß or CCL2 in response to Pam_3_CSK_4_ (Figure 8). This is consistent with the lack of effect of TAK981-mediated SUMO inhibition on the expression of these genes in response to Pam_3_CSK_4_. Together, these results demonstrate that formyl-Warnericin RK-induced deSUMOylation potentiates TNFα-induced inflammatory responses in intestinal cells.

**Figure 8:**
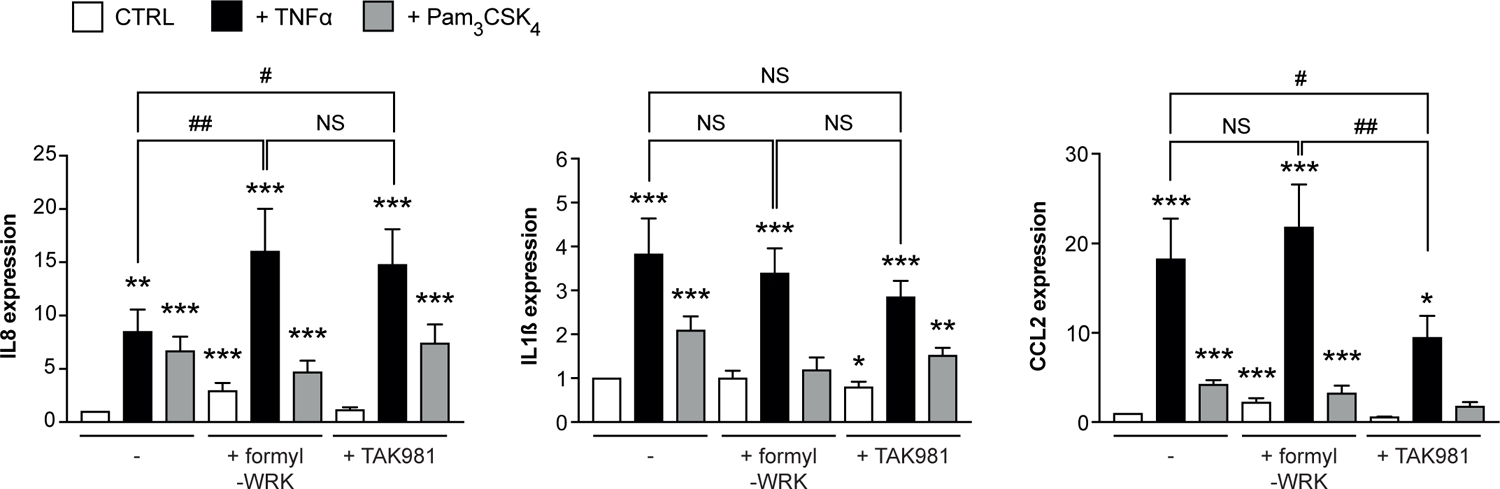
Warnericin RK potentiates TNF**α**-dependent inflammatory responses in Caco2 cells. Quantification of IL8, IL1ß and CCL2 mRNA levels in Caco2 cells pre-treated or not with formyl-WRK for 1h, or TAK981 for 2 h, and then incubated for 4 h with 100 ng/mL TNFα or 1 µg/mL Pam_3_CSK_4_. Values are expressed as fold change versus untreated cells (mean ± s.d.; n=4-5; *, *P*<0.05; **, *P*<0.01; ***, *P*<0.001 versus untreated cells; One-way ANOVA, with Dunnett’s correction; #, *P*<0.05; ##, *P*<0.01; NS, not significant; One-way ANOVA, with Tukey’s correction).

## Discussion

SUMOylation is a post-translational modification playing key roles in the maintenance of intestinal epithelium integrity and in the regulation of inflammatory responses^4,12,14^. Several gut bacterial pathogens were shown to interfere with host intestinal SUMOylation^17,18^. These pathogens use various mechanisms to dampen SUMO conjugation. Most of them directly target the SUMOylation machinery (either the E1, E2 or E3 SUMO enzymes) but bacterial virulence effectors with deSUMOylase activities have also been reported^17,18^. In contrast to pathogenic bacteria, mutualistic bacteria from the gut microbiota promote intestinal SUMOylation, via, for example, the production of SCFAs/BCFAs^4^. These bacterial metabolites, which are abundant in the gut, inactivate intestinal deSUMOylases and promote hyperSUMOylation of nuclear proteins^4^. Here, we show that *S. warneri*, a non-pathogenic bacterium from the human gut microbiota, has a similar impact on host SUMOylation than pathogens, as it dampens intestinal SUMOylation. *S.warneri*-induced decrease in SUMO-conjugated proteins is associated with a decrease in the level of E1 and E2 SUMO enzymes. As described above, the SUMOylation level of proteins results from the dynamic balance between SUMO-conjugation and SUMO-deconjugation reactions. Inhibition of SUMO-conjugation reactions shifts this equilibrium towards the deSUMOylated forms of proteins, due to the high activity of cellular deSUMOylases. The set of SUMO-conjugated proteins affected by this process would thus depend on their sensitivity/accessibility to deSUMOylases.

We identified Warnericin RK as the bacterial effector encoded by *S. warneri* responsible for this decrease in SUMO-conjugated proteins. This 22 amino acids-long peptide was initially identified for its antimicrobial activity against *Legionella pneumophila*^29^. It has been proposed that Warnericin RK kills *Legionella* by acting as a detergent on bacterial membranes. We show that, in addition to its antimicrobial activity, Warnericin RK also acts on eukaryotic cells. This versatility of activities has been observed in many other examples of antimicrobial peptides^32^.

The mechanism by which Warnericin RK forms pores or destabilizes eukaryotic membranes remains to be characterized. We show that formylated Warnericin RK has a higher hemolytic activity than N-terminal free Warnericin RK, as well as a more potent effect on host SUMOylation (Fig. 5). It has been proposed that formylation blocks the N-terminal amine group of Warnericin RK which thus cannot be protonated, thereby decreasing its net positive charge and increasing its hydrophobicity^33^. This increase in hydrophobicity probably facilitates the destabilization of eukaryotic membranes and the downstream induction of host deSUMOylation. Interestingly, we show that delta-hemolysin II, another hemolytic toxin produced by *S. warneri,* and also encoded by *S. epidermidis,* has no impact on host SUMOylation. This lack of effect might be due to the lower hemolytic activity of this toxin (Fig. 5).

Interestingly, the effect of Warnericin RK on host SUMOylation is quite similar to that of Listeriolysin O (LLO), a pore-forming toxin secreted by *L. monocytogenes*. In the case of LLO, pores formed by the toxin after binding to host membranes triggers an uncharacterized signalling cascade leading to the degradation of Ubc9 and to a decrease in SUMO-conjugated proteins^19^. Proteins deSUMOylated in response to LLO were mainly shown to be transcription factors^20^. Whether Warnericin RK triggers the same signalling cascade than LLO and whether deSUMOylated targets are similar between Warnericin RK and LLO remain to be determined.

In addition to its impact on host SUMOylation, *S. warneri* shares other similarities with *bona fide* intestinal pathogens, since this bacterium is able to get internalized into human intestinal cells^24,34^. Our work now paves the way for future studies to determine whether formyl-Warnericin RK is involved in *S. warneri* internalization, whether deSUMOylations triggered by *S. warneri* have an impact on the intracellular fate of the bacterium and whether it contributes to the consequences of *S. warneri* internalization on intestinal cell responses.

Our results indicate that the deSUMOylations triggered by Warnericin RK promote inflammatory responses in the intestinal epithelium. Interestingly, previous studies showed that deSUMOylation of specific host proteins, such as PML, could be sensed as a danger signals by host cells and trigger anti-bacterial responses^27^. More generally, since many transcription factors are regulated by SUMOylation, alteration of SUMO-conjugation may change the activity of these factors and thus participate to transcription remodelling^35^.

We identify here an example of gut bacteria dampening SUMOylation and promoting inflammation. This contrasts with the previously identified SCFA/BCFA-producing bacteria that increase SUMOylation and dampen inflammation. Our work thus unveils a balance in the gut between bacterial signals promoting or dampening SUMOylation. Changes in the composition of the gut microbiota, for example in the case of a dysbiosis, may thus alter this balance, with potential important repercussions on host intestinal physiology. Interestingly, deregulation of SUMOylation has been implicated in inflammatory diseases such as Inflammatory Bowel Diseases (IBD). Indeed, patients with Crohn’s disease or Ulcerative colitis show a decrease in colonic SUMOylation associated with a decrease in Ubc9 level^13^. This decrease in Ubc9 level was also observed in a mouse model of colitis^13^. By showing that deSUMOylation triggered by *S. warneri* promotes inflammatory responses, our results suggest that an increase in the abundance of *S. warneri* in the gut (or, more generally, an increase of bacteria inhibiting SUMOylation) may promote the onset or maintenance of inflammation and thus participate to intestinal inflammatory diseases.

In conclusion, our results unveil for the first time that non-pathogenic gut bacteria such as *Staphylococcus warneri* modulate intestinal cell activity by dampening SUMOylation. This finding strengthens the importance of SUMOylation in intestinal homeostasis and open new research avenues to determine how disequilibrium between pro- and anti-SUMO gut bacterial effectors may participate to human intestinal pathologies.

## Acknowledgments

This work was supported by INSERM, Rouen University, Janssen Horizon, the European Union, the Regional Council of Normandie (RIN Doctorant) and the french National Research Agency (ANR) under grant “SUMONING ANR-22-CE14-0064-01”. Europe gets involved in Normandie with European Regional Development Fund (ERDF).

## Data availability statement

The authors confirm that the data supporting the findings of this study are available within the article and its supplementary materials.

---

**Figure S1:** Viability of HeLa and Caco2 cells incubated with formyl-WRK. Viability of HeLa (A) or Caco2 (B) cells incubated for 1h, 4h or 5h with increasing concentrations of formyl-WRK (mean ± s.d.; n=4; *, *P*<0.05; **, *P*<0.01; ***, *P*<0.001; One-way ANOVA, with Dunnett’s correction).

**Figure S2:** Lack of effect of Delta-lysin II on SUMOylation in HeLa cells. Quantification of SUMO2/3-conjugated proteins (above 50 kDa), SAE1, SAE2 and Ubc9 levels, after normalization by actin levels, in HeLa cells treated with HldII or formyl-HldII for 1 h. Values are expressed as fold-change versus untreated cells (mean ± s.d.; n=4-5; *, *P*<0.05; One-way ANOVA, with Dunnett’s correction).

