## Supplementary figures and images for "*Staphylococcus warneri* dampens SUMOylation and promotes intestinal inflammation"

### Supplemental Figure 1

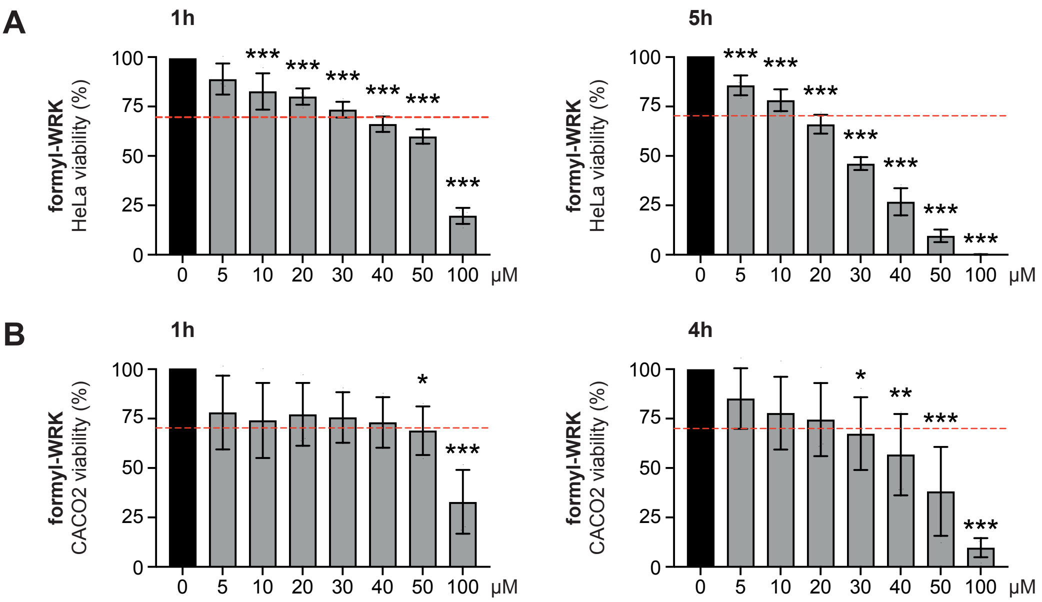


#### SUPPLEMENTARY FIGURE 1

### Supplemental Figure 2

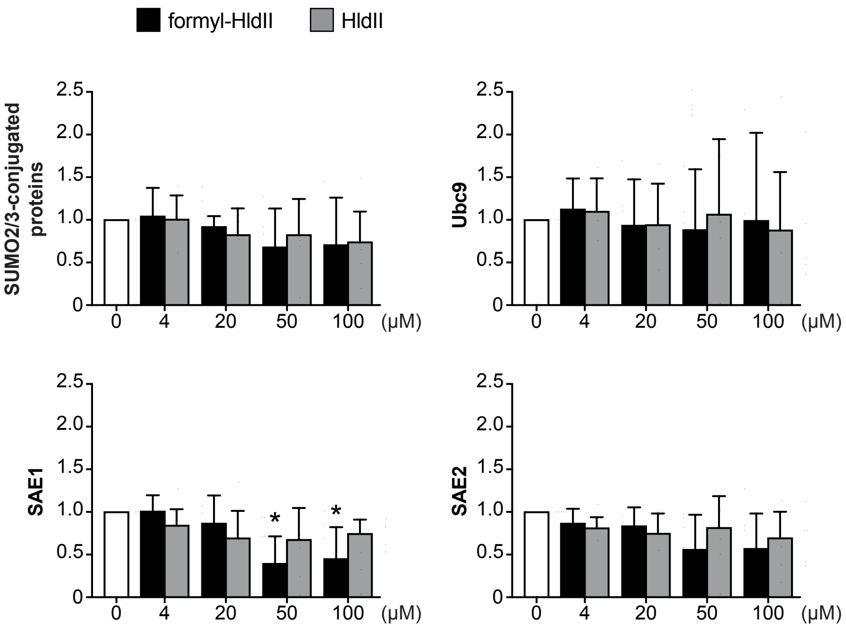


#### SUPPLEMENTARY FIGURE 2
